# Impact of different amino acid substitutions in penicillin-binding protein 3 on beta-lactam susceptibility in *Haemophilus influenzae*

**DOI:** 10.1101/846428

**Authors:** Josiane Reist, Janina Linnik, Urs Schibli, Adrian Egli, Vladimira Hinić

**Affiliations:** Division of Clinical Bacteriology and Mycology, University Hospital Basel, Basel, Switzerland; Applied Microbiology Research, Department of Biomedicine, University of Basel, Basel, Switzerland; Department of Biosystems Science and Engineering, ETH Zurich, Basel, Switzerland; Swiss Institute for Bioinformatics, Basel, Switzerland; Bakteriologisches Institut Olten, Kantonsspital Olten, Switzerland

**Author notes:** These first authors contributed equally to this article (Josiane Reist, Janina Linnik). These senior authors contributed equally to this article (Adrian Egli, Vladimira Hinić). Corresponding author. Mailing address: Division of Clinical Microbiology, University Hospital Basel, Petersgraben 4, 4031 Basel, Switzerland. Parts of this study were presented at the European Congress of Clinical Microbiology and Infectious Diseases (ECCMID), held in Vienna from 22 to 25 April 2017.

**Keywords:** *Haemophilus influenzae*, beta-lactam antimicrobials, beta-lactam resistance, PBP3, genotype, phenotype

## Abstract

**Purpose:** Beta-lactam antibiotics in combination with a beta-lactamase inhibitor are the first-line treatment option for *Haemophilus influenzae* infections. However, beta-lactamase-independent resistance to beta-lactams is increasing. This resistance mechanism has been linked to amino acid substitutions in the penicillin-binding protein 3 (PBP3), but how these substitutions lead to decreased binding affinities to certain beta-lactam antimicrobials remains unknown.

**Methods:** We investigated beta-lactam resistance and amino acid substitutions in PBP3 from fifty-three clinical isolates of *H. influenzae* collected in Switzerland from January to April 2016. Identification of key polymorphisms and classification of strains into PBP3 amino acid substitution groups I, II, and M was done as previously described. Based on published PBP3 crystal structures, we investigated how the group-specific amino acid substitutions impact the beta-lactam binding site.

**Results:** We found that both group I and group II substitutions disrupt the Asn526-Arg517-Glu324 interaction, which might affect the configuration of the beta-lactam binding site. Amino acid substitutions in group M strains are distant from the active site and have most likely no impact on beta-lactam binding. In accordance with this observation, all group M strains showed minimal inhibitory concentrations (MICs) within the susceptible range for all tested antimicrobials and were not significantly different to the wild type (beta-lactamase producers excluded), while group I and group II strains showed significantly higher MICs for beta-lactam antimicrobials.

**Conclusion:** Group M strains are phenotypically equal to the wild type, while amino acid substitutions of group I and group II might affect the beta-lactam binding through a common mechanism by disrupting the Asn526-Arg517-Glu324 interaction.

## Introduction

Beta-lactam antibiotics are considered to be the first-line treatment option for *Haemophilus influenzae* infections: aminopenicillins used in combination with a beta-lactamase inhibitor for respiratory and other non-invasive infections; and third-generation cephalosporins for invasive infections such as meningitis (1). Although the percentages of beta-lactamase positive isolates in European countries are relatively low (Turkey 6.8%, Germany 9.3%, France 11.9%, Poland 17.9%) (2–5), the percentages in some Asian countries are much higher (China 31.0%, Vietnam 40.5%) (6, 7). There are two main beta-lactamase types found in *H. influenzae*: TEM-1 and ROB-1 (8). According to several studies, ROB-1 beta-lactamase is very rare and mostly found in North America, while TEM-1 type is globally distributed (8–10). These beta-lactamases confer resistance to aminopenicillins and are easily detected by using a simple nitrocefin-based beta-lactamase test.

However, the appearance and spreading of beta-lactamase-independent beta-lactam resistance is more worrying (11, 12). The beta-lactam resistence in these isolates is mostly due to amino acid substitutions in the *ftsI* gene which lead to alterations in penicillin-binding protein 3 (PBP3) and consequently a decreased affinity for binding of certain beta-lactam antibiotics (13, 14). Based on the beta-lactamase production and presence of *ftsI* mutations, the strains can be classified into four susceptibility groups (**Table 1a**) as previously described by Ubukata et al., Dabernat et al. and García-Cobos et al. (15–17). The strains with low-level PBP3-mediated resistance such as group I and II (low-rPBP3) show ampicillin minimal inhibitory concentrations (MICs) between 0.5 and 2 mg/L, whereas strains with high-level resistance (high-rPBP3) such as group III exhibit higher ampicillin MICs between 1 and 16 mg/L, often combined with reduced susceptibility to cephalosporins (11, 17–22). Although group III isolates are mainly distributed in some Asian countries such as Japan and Korea (23, 24), these more resistant strains, especially group III-like have been detected on rare ocasions in some European countries (3, 14, 20, 25, 26).

**Table 1.** **(a)** Categorization of *H. influenzae* strains into four aminopenicillin susceptibility groups based on the beta-lactamase production and presence of *ftsI* mutations. **(b)** PBP3 mutation groups and corresponding key amino acid substitutions.

|  |  |  |
| --- | --- | --- |
| (a) | Abbreviation | Genotype |
|  | gBLNAS | Beta-lactamase negative and ampicillin susceptible without resistance mechanism |
|  | gBLNAR | Beta-lactamase negative and ampicillin resistant with PBP3 mutations |
|  | gBLPAR | Beta-lactamase positive and ampicillin resistant without PBP3 mutations |
|  | gBLPACR | Beta-lactamase positive and amoxicillin-clavulanic acid resistant with PBP3 mutations |
| (b) | PBP3 mutation group | Key amino acid substitution |
|  | I<sup>1</sup> | Arg517His |
|  | II<sup>1</sup> | Asn526Lys |
|  | IIa<sup>2</sup> | Asn526Lys (the only substitution) |
|  | IIb<sup>2</sup> | Asn526Lys, Ala502Val |
|  | IIc<sup>2</sup> | Asn526Lys, Ala502Thr |
|  | IId<sup>2</sup> | Asn526Lys, Ile449Val |
|  | III<sup>1</sup> | Met377Ile, Ser385Thr, Leu389Phe, Asn526Lys |
|  | III-like<sup>3</sup> | Group III mutations without Asn526Lys or with additional combinations at the KTG or STVK motif |
|  | M<sup>3</sup> | Miscellaneous |

<sup>1</sup>According to Ubukata et al. (15)
<sup>2</sup>According to Dabernat et al. (16)
<sup>3</sup>According to Garcia-Cobos et al. (17)

PBPs are transpeptidases or carboxypeptidases involved in peptidoglycan metabolism. They harbour three specific motifs: SXXK, (S/Y)XN and (K/H) (S/T)G, which define the active site of the serine penicillin-recognizing enzyme family (15, 27). The serine in the SXXK motif is crucial to the catalytic mechanism (27).

The aim of our study was to investigate the impact of PBP3 amino acid substitutions in clinical isolates of *H. influenzae* on (i) antimicrobial susceptibility, (ii) the beta-lactam binding site based on published PBP3 crystal structures, and (iii) to investigate, whether there is an association between antimicrobial susceptibility and the location of the substituted amino acids.

## Materials and methods

### Clinical specimens and reference strains

The clinical isolates of *H. influenzae* were collected prospectively from January to April 2016 from in-patients of the University Hospital Basel, Switzerland. Three isolates originated from patients of the Cantonal Hospital in Olten, Switzerland. One isolate originated from the UK NEQAS external quality assessment services, Scotland. Only one sample per patient was included in the study. Thirty-four samples were obtained from respiratory samples: sputum (n=20), tracheal secretion (n=8), bronchial secretion (n=4), and bronchoalveolar lavage (n=2). Nineteen samples were collected from other body sites: conjunctiva (n=8), ear canal (n=3), cerebrospinal fluid (n=2) and others (n=6).

### Isolation and storage of the isolates

The isolation of *H. influenzae* from clinical samples was carried out by standard protocols including Haemophilus Chocolate 2 agar (HAE2) or chocolate PolyViteX (PVX) agar (bioMérieux, Marcy-l’Étoile, France) incubated at 36 °C under the atmosphere containing 5% CO_2_. Colonies were identified with matrix-assisted laser desorption ionization time of flight mass spectrometry (MALDI-TOF MS; Bruker, Bremen, Germany; MBT 6903 MSP Library version). All *H. influenzae* isolates were frozen at −70 °C in cryogenic Microbank^TM^ vials (Pro-Lab Diagnostics, Birkenhead, UK). Prior to testing, the strains were cultured on PVX agar with subculture after 24 hours.

### Antimicrobial susceptibility testing

Minimal inhibitory concentrations (MICs) determination was performed with Etest® (bioMérieux). The following antimicrobials were tested: ampicillin, amoxicillin, amoxicillin-clavulanic acid, piperacillin, piperacillin-tazobactam, cefuroxime, and meropenem. The MIC breakpoints and screening results for beta-lactam resistance with 1 U benzylpenicillin disc were interpreted as defined by the European Committee on Antimicrobial Susceptibility Testing (EUCAST, version 7.1) (28), except for piperacillin-tazobactam which was interpreted as defined by the Clinical and Laboratory Standards Institute (CLSI, 27th edition) (29) due to no existing EUCAST interpretation criteria. Piperacillin was interpreted according to CLSI guidelines for piperacillin-tazobactam, because there are no interpretation criteria in either EUCAST or CLSI guidelines for this antimicrobial.

### Statistical analysis of MICs for comparison of strains with amino acid substitutions to wild type strains

Wilcoxon rank-sum test was used to compare MICs between strains as follows: For all tested antimicrobials, we compared the MICs of group IIa, IIb, IId and group M strains to the respective MICs of the wildtype. Group I strains were excluded from the analysis, because we found only two strains in this group. Since we were interested in beta-lactamase independent resistance, all beta-lactamase producing strains were excluded as well. To correct the obtained p-values for multiple hypothesis testing, we used the Benjamini-Hochberg-Yekutiely procedure that controls the false discovery rate (30). The analysis has been done in R. All analysis scripts including raw data and figure-generating scripts are available on gitlab: https://gitlab.com/csb.ethz/haemophilus-pbp3-manuscript (online ressource **ESM1)**.

### Phenotypic and genotypic determination of beta-lactamase production

Beta-lactamase production was determined with BBL^TM^ Cefinase^TM^ Paper Discs (Beckton Dickinson, Franklin Lakes, NJ, USA) according to manufacturer’s instructions. Presence of TEM-1 beta-lactamase encoding gene was determined by conventional PCR with primers described by Dabernat et al. (16). The following reference strains were used as controls for TEM-1 PCR: *H. influenzae* ATCC 49247 (BLNAR) and *H. influenzae* ATCC 35056 (BLPAR).

### Determination of *ftsI* mutations and identification of PBP3 amino acid substitution groups

Nucleic acid extraction was performed with Advanced XL EZ1 (Qiagen, Hilden, Germany). The digestion step using proteinase K was performed for 10 minutes at 56 °C and 10 minutes at 95 °C followed by purification with EZ1 Tissue card according to the manufacturer’s protocol. DNA extracts were eluted in 100 μl elution buffer. Amplification of *ftsI* gene was performed by using the conventional PCR assay described by Cerquetti et al. (31). PCR products were purified with ExoSAP-IT® (USB Corporation, Cleveland, OH, USA) purification kit. The sequencing of the PCR amplicon was performed by Microsynth AG (Balgach, Switzerland) using Sanger sequencing (sequencing primer^frw^: 5’-GCGGATAAAGAACGAATTGC-3’ (14), sequencing primer^rev^: 5’-CTGGATAATTCTGTCTCAGA-3’ (31)). Sequences were aligned with Lasergene SeqMan Pro (DNASTAR, Madison, WI, USA) and translated into amino acid sequences using the ExPASy translate tool (http://web.expasy.org/translate/). To identify amino acid substitutions, the translated amino acid sequences were aligned using Clustal Omega (http://www.ebi.ac.uk/Tools/msa/clustalo/) to reference strain *H. influenzae* RD KW20 (Acc. No. Genbank NC_000907). Key amino acid changes were grouped as reported previously, see **Table 1b** (15–17).

### Investigation of group-specific mutations in the PBP3 crystal structure

At the time of this study, there were no crystal structures of PBP3 from *Haemophilus* spp. in the Worldwide Protein Data Bank (wwPDB; www.wwpdb.org). Therefore, we selected structures from other gram-negative rods *Escherichia coli* (PDB ID 4BJP, 2.5 Å resolution) and *Pseudomonas aeruginosa* (PDB ID 3PBR, 1.95 Å resolution) for our analysis. Fortunately, the amino acid sequences of the three conserved PBP3 domains of *E. coli* and *P. aeruginosa* are identical to those of *H. influenzae*, except for one conservative amino acid substitution in *P. aeruginosa*, where serine is replaced by threonine in the (K/H) (S/T)G motif. In addition, the root mean square deviation (RMSD) of all matched atoms in the PBP3 crystal structures from *E. coli* and *P. aeruginosa* is 1.166 Å, which stands for high structural similarity (32). We used Clustal Omega to align the PBP3 amino acid sequences and to determine the sequence identity between the different species, i.e. *H. influenzae* RD KW20 (UniProt Entry P45059), *Escherichia coli* (GI:635575685), and *Pseudomonas aeruginosa* (UniProt Entry Q51504). Crystal structures of PBP3 from *E. coli* and *P. aeruginosa* were aligned with PyMOL (The PyMOL Molecular Graphics System, Version 2.0 Schrödinger, LLC). To investigate whether specific amino acid substitutions impact the meropenem binding site, we constructed three-dimensonal models of amino acid substitutions in *H. influenzae* based on the *P. aeruginosa* crystal structure (PDB ID code 3PBR).

## Results

### Identification of PBP3 amino acid substitution groups

Based on the PBP3 amino acid sequence deduced from the *ftsI* gene sequence, we assigned the investigated strains (n=53) to following groups as previously described (**Table 1b**): wild type (n=17), group I (n=2), group II (a, b and d; n=21) or group M (miscellaneous, n=13) (**Table 2**). No isolates belonging to groups IIc, III or III-like were found. Eight out of 13 group M isolates with three different mutation patterns (Asp350Asn and Val547Ile; Val547Ile; Asp350Asn and Val547Ile and Asn569Ser) were described previously by Cherkaoui et al. (14, 33). The remaining substitution combinations in group M have been identified for the first time in four strains of this study (**Table 2**).

**Table 2.**
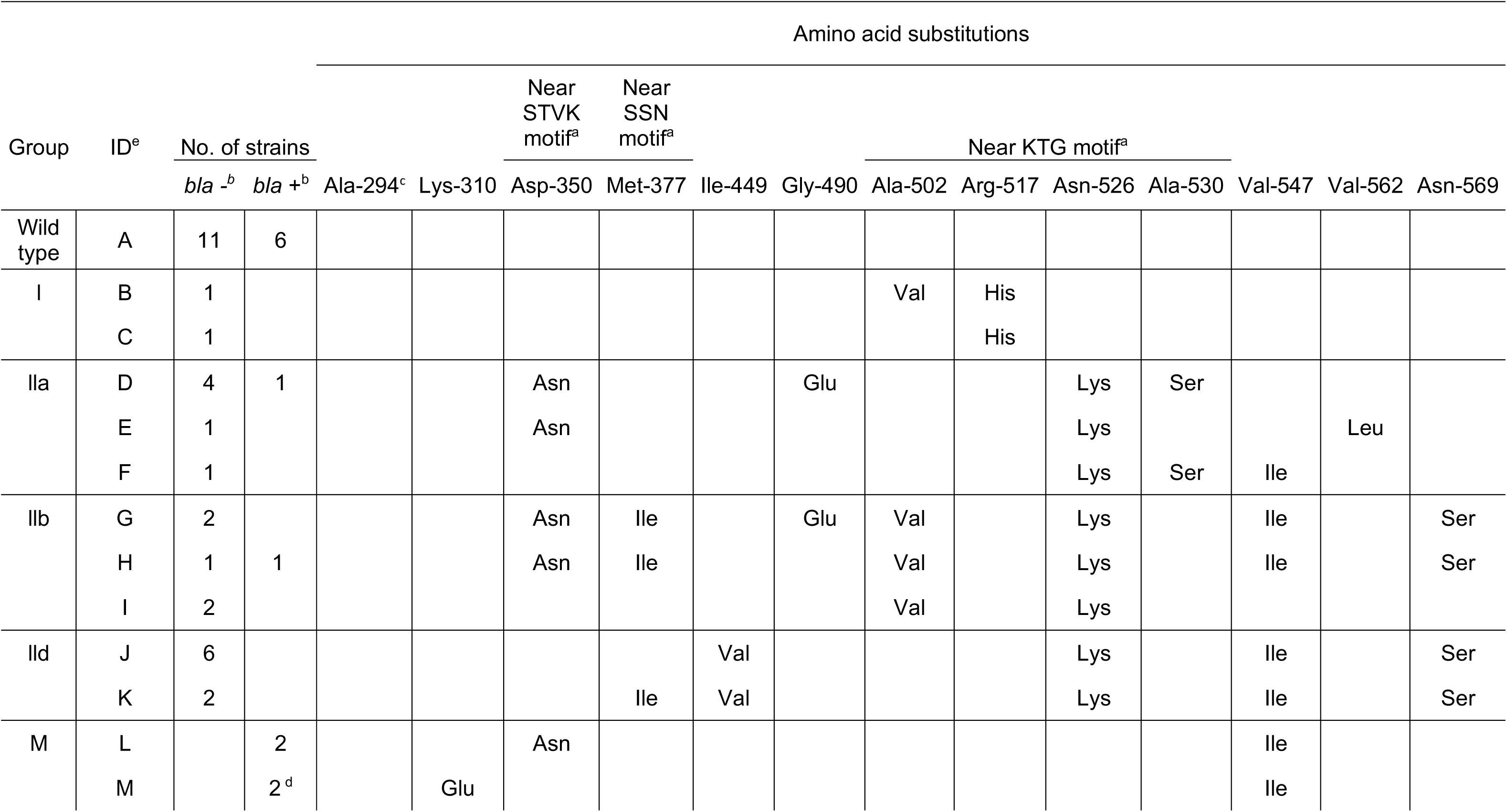
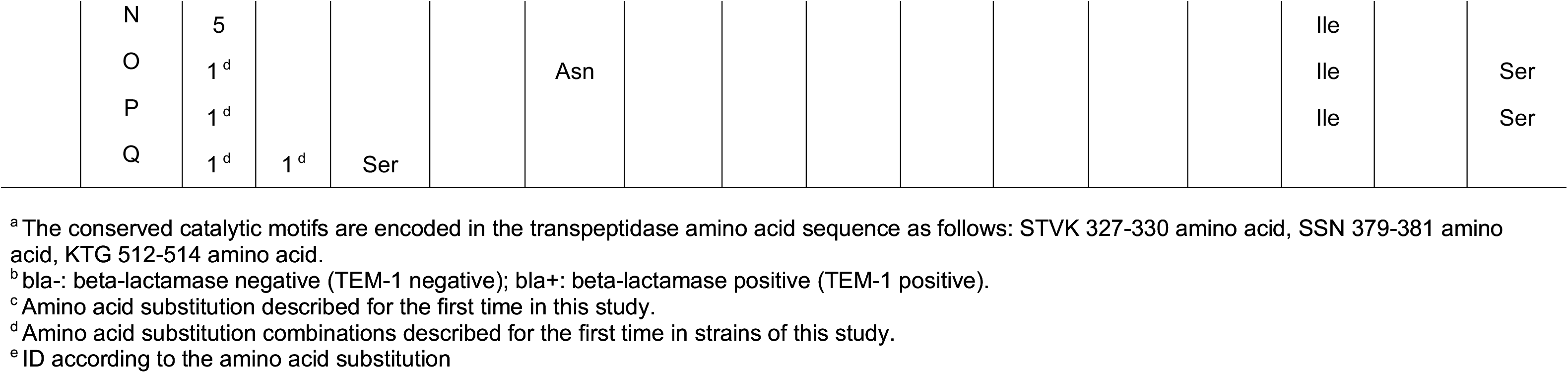
Amino acid substitutions in the transpeptidase domain of the *ftsI* gene found in the strains of this study (n=53).

### MICs for beta-lactam antibiotics of different PBP3 amino acid substitution groups

All determined MICs are summarized in **Table 3** and shown in **Figure 1**. All strains with amino acid substitutions in PBP3 belonging to groups I, II and M (beta-lactamase producers excluded) showed very low MICs for piperacillin and piperacillin-tazobactam (range 0.016-0.25 and <0.016-0.25 mg/l, respectively), which was not observed for any other penicillin with or without beta-lactamase inhibitor (**Table 3 and Figure S1**). For all tested antimicrobials, we compared the MICs of group IIa, IIb, IId and group M isolates (beta-lactamse producers excluded) to the respective MICs of the wildtype. Group I isolates have been excluded due to low sample size (n=2). We found no significant difference between group M and the wild type isolates. In contrast, all group II isolates showed significantly higher MICs (p-values ranging from 0.007 to 0.016, see **Figure 1**) except for piperacillin and piperacillin-tazobactam.

**Fig. 1.**
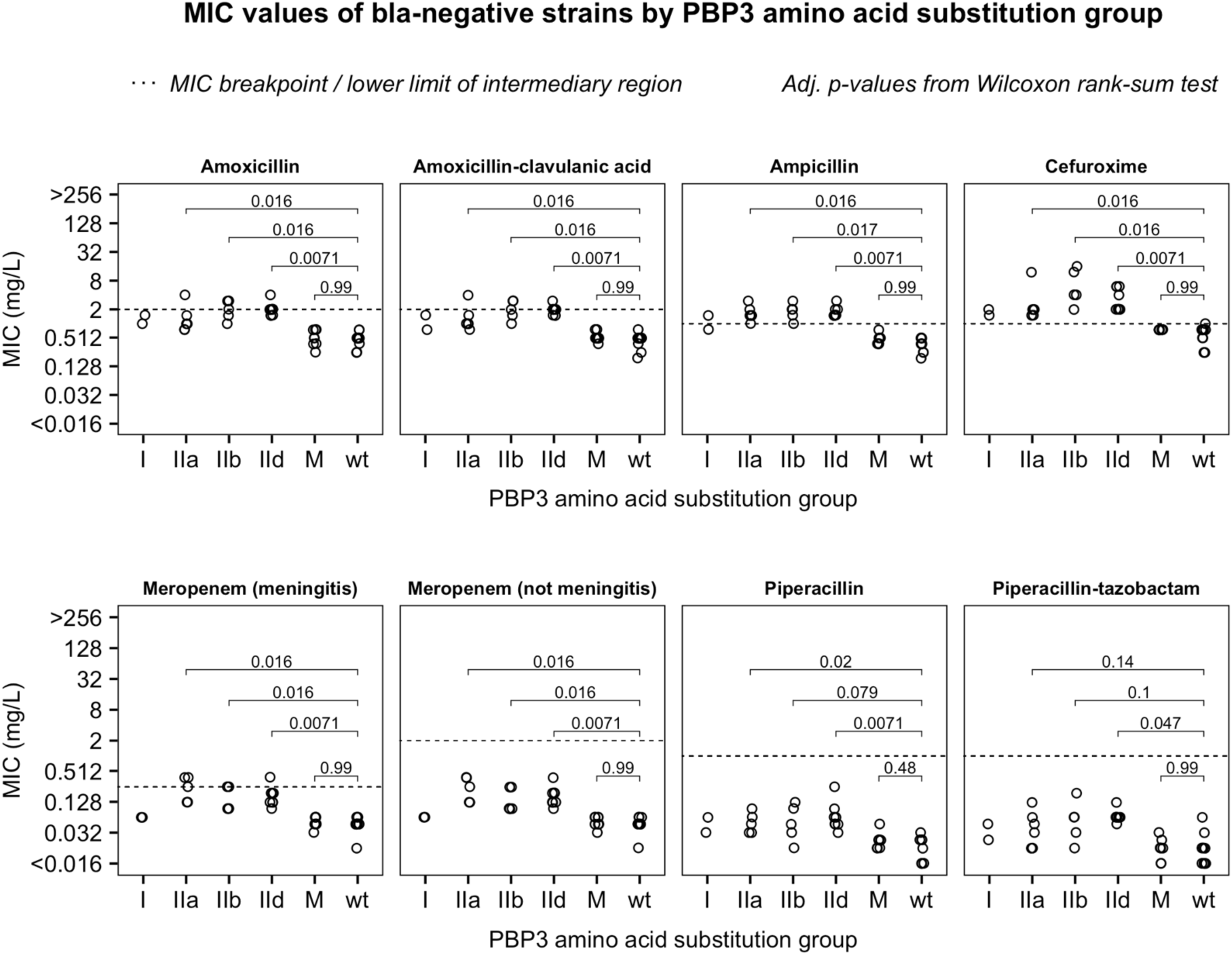
MIC values of beta-lactamase negative strains. Wilcoxon rank-sum test was used to compare MIC values of strains with amino acid substitutions to the wildtype. The numbers show the adjusted p-values (Benjamini-Hochberg-Yekutiely correction). The dotted line indicates the MIC breakpoint or, if applicable, the lower limit of the intermediary region. All antibiotic breakpoints are as defined by EUCAST with the exception of piperacillin and piperacillin-tazobactam which are according to CLSI.

**Table 3.** Results of minimal inhibitory concentration (MIC) testing according to amino acid substitution groups.

| Group | ID<sup>g</sup> | No. of isolates |  | Ampicillin | Amoxicillin | Amoxicillin-clavulanic<br>acid | Piperacillin | Piperacillin-tazobactam | Cefuroxime<br>(iv<sup>c</sup>) | Meropenem<br>(infections<br>other than<br>meningitis) | Meropenem<br>(meningitis) |
| --- | --- | --- | --- | --- | --- | --- | --- | --- | --- | --- | --- |
|  |  | *bla* -<sup>b</sup> | *bla* +<sup>b</sup> |  |  |  |  |  |  |  |  |
| Wild<br>type | A |  | 6 | 4->256 | 3->256 | 0.38-1.5 | 2-24 | 0.016- 0.064 | 0.38-1 | 0.032-0.064 | 0.032-0.064 |
|  | A | 11 |  | 0.25-0.5 | 0.25-0.75 | 0.19-0.75 | <0.016-0.032 | <0.016-0.064 | 0.19-1 | 0.016-0.064 | 0.016-0.064 |
| I | B | 1 |  | 1.5 | 1.5 | 1.5 | 0.032 | 0.023 | 2 | 0.064 | 0.064 |
|  | C | 1 |  | 0.75 | 1 | 0.75 | 0.064 | 0.047 | 1.5 | 0.064 | 0.064 |
| IIa | D |  | 1 | >256 | >256 | 1 | 8 | 0.064 | 2 | 0.19 | 0.19 |
|  | D | 4<sup>e</sup> |  | 1- 2 | 0.75-1.5 | 0.75-1.5 | 0.032-0.094 | 0.016-0.064 | 1.5-2 | 0.125-0.38 | 0.125 -0.38 |
|  | E | 1 |  | 3 | 4 | 4 | 0.064 | 0.032 | 12 | 0.38 | 0.38 |
|  | F | 1 |  | 1.5 | 1 | 1 | n.d.<sup>d</sup> | 0.047 | 1.5 | n.d.<sup>d</sup> | n.d.<sup>d</sup> |
| IIb | G | 2 |  | 1-2 | 1- 3 | 1- 3 | 0.016-0.094 | 0.016-0.064 | 4-16 | 0.094-0.25 | 0.094-0.25 |
|  | H |  | 1 | >256 | >256 | 3 | >256 | 0.25 | 16 | 0.094 | 0.094 |
|  | H | 1 |  | 3 | 2 | 2 | 0.125 | 0.19 | 12 | 0.094 | 0.094 |
|  | I | 2 |  | 1.5-2 | 1.5- 3 | 1.5- 3 | 0.032-0.047 | 0.032-0.064 | 2- 4 | 0.094-0.25 | 0.094-0.25 |
| II d | J | 6<sup>f</sup> |  | 1.5-2 | 1.5-2 | 1.5-2 | 0.032-0.25 | <0.016-0.125 | 2- 6 | 0.094-0.19 | 0.094-0.19 |
|  | K | 2 |  | 2-3 | 2- 4 | 2- 3 | 0.064-0.094 | 0.064 | 6 | 0.19-0.38 | 0.19 -0.38 |
| M | L |  | 2 | >256 | >256 | 1.-1.5 | 2-8 | <0.016-0.016 | 1 | 0.094-0.125 | 0.094-0.125 |
|  | M |  | 2 | >256 | >256 | 1 | 8-64 | 0.016-0.023 | 0.75 | 0.064 | 0.064 |
|  | N | 5 |  | 0.38-0.5 | 0.25-0.75 | 0.38-0.75 | 0.016-0.047 | <0.016-0.032 | 0.75 | 0.032-0.094 | 0.047-0.094 |
|  | O | 1 |  | 0.75 | 0.75 | 0.75 | 0.023 | 0.016 | 0.75 | 0.064 | 0.064 |
|  | P | 1 |  | 0.5 | 0.75 | 0.75 | 0.023 | 0.016 | 0.75 | 0.064 | 0.064 |
|  | Q |  | 1 | >256 | >256 | 1 | 32 | 0.023 | 0.75 | 0.047 | 0.047 |
|  | Q | 1 |  | 0.38 | 0.38 | 0.5 | 0.016 | <0.016 | 0.75 | 0.047 | 0.047 |

<sup>a</sup> green: susceptible; yellow: intermediate; red: resistant.
<sup>b</sup> bla-: beta-lactamase negative; bla+: beta-lactamase positive.
<sup>c</sup> iv: refers to intravenous administration of cefuroxime;
<sup>d</sup> n.d.: not done.
<sup>e</sup> ampicillin: one isolate susceptible, three isolates resistant; meropenem: one isolate intermediate, three isolates susceptible
<sup>f</sup> cefuroxime (i/v): four isolates intermediate, two isolates resistant
<sup>g</sup> ID according to the amino acid substitution

In the benzylpenicillin disc screen, all wild type and group M isolates (except beta-lactamase producing strains) were categorized as beta-lactam susceptible (**Figure S2**). All group I and group II isolates were categorized as resistant in the beta-lactamase screen (zone diameter <=11mm), except for two strains with zone diameters of 12 mm (on the breakpoint) which were categorized as susceptible.

### Impact of group-specific amino acid substitutions on the PBP3 crystal structure

We investigated the position of the identified group-specific amino acid substitutions and their impact on the PBP3 binding site (**Figure 2**). Since there was no PBP3 crystal structure of *H. influenzae* available at the time of this study, we selected PBP3 crystal structures from *E. coli* and *P. aeruginosa* for our analysis, which amino acid sequences of the three conserved PBP3 domains were identical to those of *H. influenzae*, except for one conservative amino acid substitution in *P. aeruginosa*, where serine is replaced by threonine in the (K/H) (S/T)G motif. The overall PBP3 amino acid sequence identity between *H. influenzae* and *E. coli* is 54%, between *H. influenzae* and *P. aeruginosa* 39%, and between *E. coli* and *P. aeruginosa* 47%. These identities are sufficient for our analysis, because we are interested in the position of mutations in the three-dimensional structure and protein structure is more conserved than protein sequence (32, 34). Crystal structure alignment of PBP3 of *E. coli* and *P. aeruginosa* confirmed this statetement: the RMSD (Root Mean Square Deviation) of the alignment of all matched atoms is 1.166 Å (see **Figure 2a**), which stands for a good accordance (32). Because the *P. aeruginosa* crystal structure (PDB ID code 3PBR) contains a representation of the binding of beta-lactam meropenem and has a high resolution (1.95 Å), we used this structure to model the amino acid substitutions in *H. influenzae*. Interestingly, all relevant amino acids were identical between the two strains, i.e. the amino acids that we found to be directly or indirectly affected by the group-specific mutations Arg517His, Asn526Lys, and Val547Ile (**Figure S3**).

**Fig. 2.**
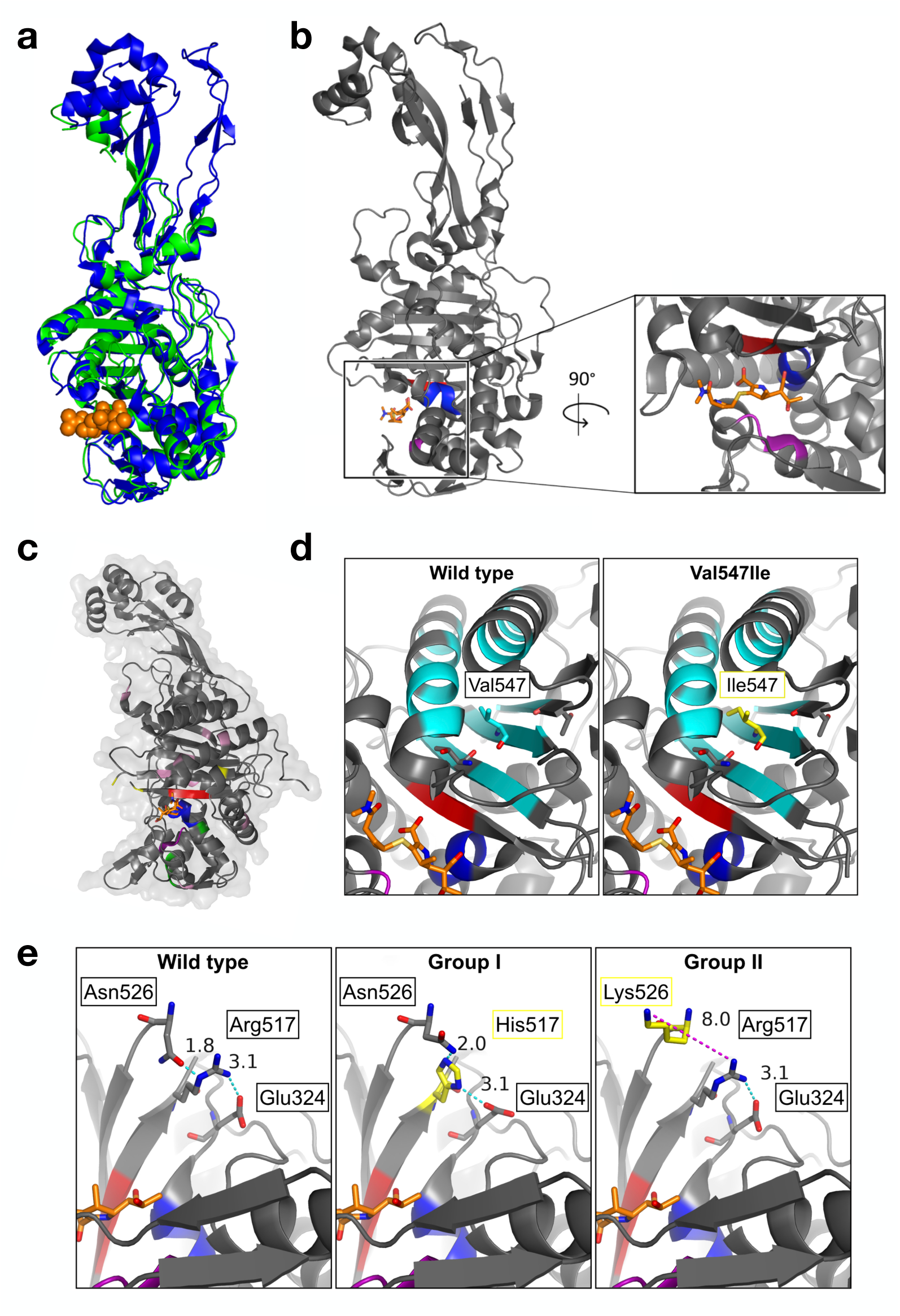
PBP3 crystal structure analysis. **(a)** Crystal structure alignment of PBP3 of *Escherichia coli* (green, PDB ID 4BJP, solved at 2.50 Å) and of *Pseudomonas aeruginosa* (blue, PDB ID 3PBR, solved at 1.97 Å). The meropenem molecule is shown in orange. The root mean square deviation of all matched atoms (2185 atoms) is 1.166 Å. **(b)** PBP3 crystal structure of *P. aeruginosa* with a molecule of meropenem (orange) bound at the active site. The transpeptidase domain belongs to the active C-terminus. The three catalytic domains are highlighted in blue (SXXK), purple ((S/Y)XN) and red ((K/H)(S/T)G motif). **(c)** Localization of amino acid substitutions of PBP3 groups in the PBP3 crystal structure of *P. aeruginosa* (PDB ID 3PBR): group I and II are highlighted in yellow, group III in green and group M in pink. The three catalytic domains are highlighted in blue (SXXK), purple ((S/Y)XN) and red ((K/H)(S/T)G motif). The meropenem molecule is shown in orange. **(d)** Graphic representation of the most common group M mutation Val547Ile (shown in yellow) in the PBP3 crystal structure of *P. aeruginosa* (PDB ID 3PBR). Val547 lies in a hydrophobic pocket where it is surrounded by aliphatic amino acids (shown in cyan), except for Thr532 and Thr281, which are polar. The substitution of an aliphatic amino acid by another aliphatic amino acid (valine to isoleucine) has most likely no impact on the protein structure. The three catalytic domains are highlighted in blue (SXXK), purple ((S/Y)XN) and red ((K/H)(S/T)G motif). The meropenem molecule is shown in orange. **(e)** Investigation of the group I mutation Arg517His and group II mutation Asn526Lys (both shown in yellow) based on the PBP3 structure of *P. aeruginosa* (PDB ID 3PBR). Attracting interactions are shown in cyan (i.e. hydrogen bonds and electrostatic attraction between Arg-Glu), repelling interactions in pink (electrostatic repulsion between Lys-Arg). Numbers indicate the distance between atoms in Ångström. Both mutations are able to disrupt the Arg517-His517-Glu324 interaction. The three catalytic domains are highlighted in blue (SXXK), purple ((S/Y)XN) and red ((K/H)(S/T)G motif). The meropenem molecule is shown in orange.

**Figure 2b** shows the location of the conserved catalytic domains of the beta-lactam binding site. None of the found mutations were directly located at the binding site but all substitutions of group M were distant from the beta-lactam binding site, whereas group I and II substitutions were closer to the binding site (**Figure 2c**).

The amino acid substitution Val547Ile were found in almost all group M strains and some group II strains (see **Table 2**). This conservative mutation has most likely no impact on protein structure: Val547 is located in a hydrophobic pocket where it is surrounded by aliphatic amino acids except for Thr532 and Thr281 (**Figure 2d**) the substitution of the aliphatic amino acid valine by the aliphatic amino acid isoleucine has most likely no influence on the protein structure.

The amino acid substitution Arg517His occured in group I strains, and the substitution Asn526Lys occurred in group II strains. These substitutions are adjacent and lie in neighboring beta turns of the beta sheet that harbours the KTG motif. We speculate that Arg517 and Asn526 are part of an important Arg517-Asn526-Glu324 interaction (**Figure 2e**, left panel) for the following reasons: (a) Arg517 and Asn526 are able to build a hydrogen bond and to connect the two beta turns, thus stabilizing this important region; (b) Arg517 is able to build a hydrogen bond to Glu324, thus connecting the alpha helix that harbours the STVK motif to the beta sheet that harbors the KTG motif. In addition, the positive charge of arginine and the negative charge of glutamate attract each other. These interactions between Arg517, Asn526 and Glu324 could stabilize the configuration of the two functional motifs KTG and STVK.

As shown in the middle panel of **Figure 2e**, the group I mutation Arg517His results in the loss of the electrostatic attraction to Glu324; His517 is able to replace Arg517 as a proton acceptor in the hydrogen bond to Asn526, and can serve as a proton donor in the hydrogen bond to Glu324. Yet, this substition might impact the protein structure, because (a) hydrogen bonds are weaker than electrostatic interactions, and (b) histidine is less flexible than arginine, which means that the Arg517-His517-Glu324 interaction has less degrees of freedom than the Arg517-Asn526-Glu324 bridge.

As shown in the right panel of **Figure 2e**, the group II mutation Asn526Lys results in the loss of the hydrogen bond to Arg517. In addition, lysine is negatively charged such as arginine, which leads to an electrostatic repulsion. As a result, the two beta turns might now be repelled from each other instead of attracted.

### Genotypic and phenotypic detection of beta-lactamase production

In 25% of the isolates tested (13/53), beta-lactamase was detected by Cefinase^TM^ test. The same 13 strains were also found positive with TEM-1 PCR, and the remaining 40 strains were TEM-1 negative (**Table 2**).

### Susceptibility according to four aminopenicillin susceptibility groups

Based on the MIC distributions, results of benzylpenicillin disc screen and conclusions drawn from the crystal structure binding sites, group M isolates were considered to belong either to gBLNAS or gBLPAR, while group I and II isolates were considered to belong to gBLNAR or gBLPACR. The classification of the investigated strains according to beta-lactamase production and amino acid substitution groups is as follows: 36% (19/53) gBLNAS, 39% (21/53) gBLNAR, 21% (11/53) gBLPAR and 4% (2/53) gBLPACR. The results of MIC testing according to four aminopenicillin susceptibility groups is shown in **Table 4**: 81% (17/21) of gBLNAR strains were categorized as amoxicillin and amoxicillin-clavulanic acid susceptible and 9.5% (2/21) as ampicillin susceptible. With EUCAST breakpoints for *H. influenzae* infections other than meningitis, all gBLNAR strains were meropenem susceptible, while when breakpoints for meningitis were applied, 15% (3/20) were categorized as intermediate (**Table 4**). The number of gBLPACR strains in the study was only two, and therefore no reliable classification according to susceptibility phenotype could be made for this group.

**Table 4.**
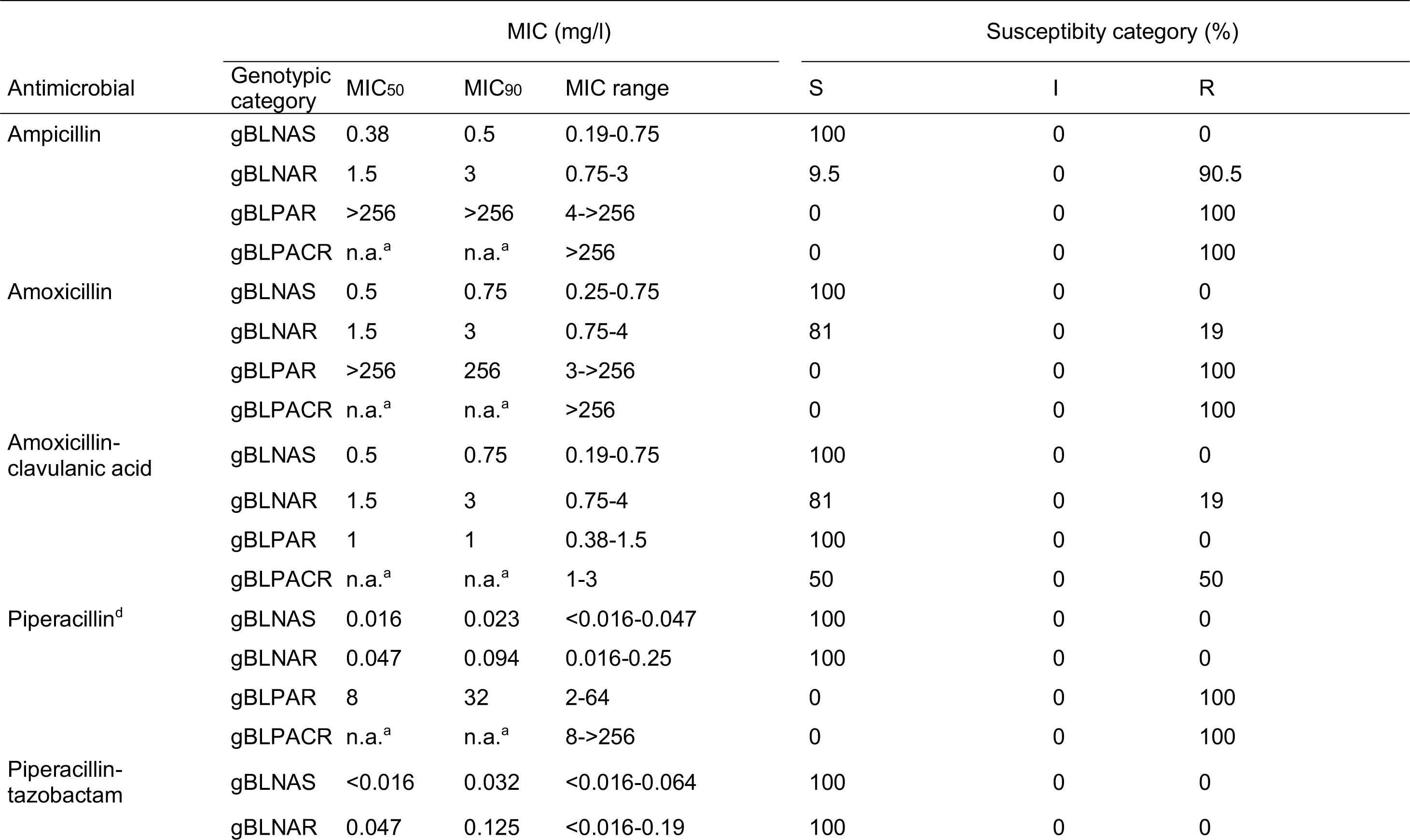
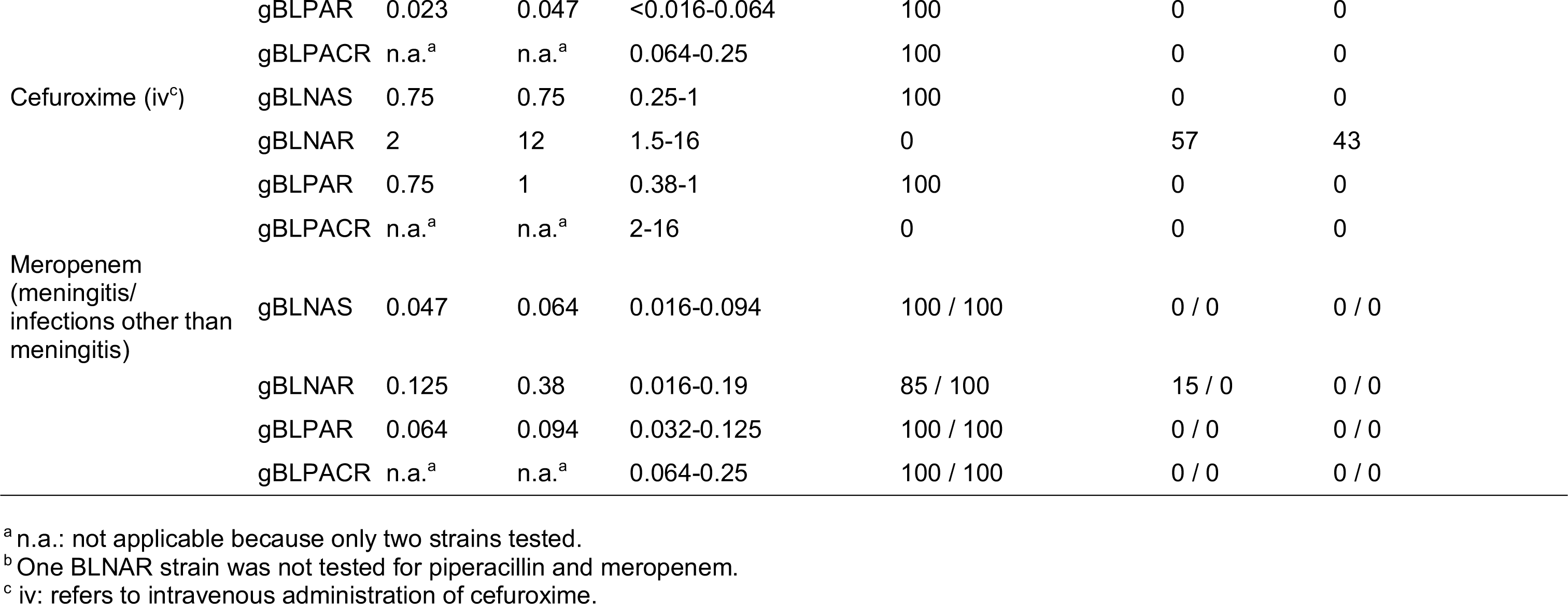
Results of MIC testing according to four aminopenicillin susceptibility groups (gBLNAS n=19, gBLNAR n=21, gBLPAR n=11, gBLPACR

## Discussion

In recent years, alterations in PBP3 leading to decreased binding affinity of certain beta-lactam antibiotics have emerged as a prevalent resistance mechanism in *H. influenzae* (35). In this study, we investigated the mutations in the *ftsI* gene encoding for PBP3 within a collection of clinical isolates from two Swiss hospitals. To our knowledge, our study investigated for the first time the location and potential impact of the polymorphisms in the PBP3 crystal structure of clinical isolates of *H. influenzae* with a detailed description of amino acid interactions. The most frequently found group of amino acid substitutions belonged to group II (19/53; 35.8%), which was also observed in studies from other European countries (3, 14, 20, 21, 36). We could not detect strains of the group III, which harbours high-level PBP3 alterations (11, 35).

All wild type strains showed MICs within the susceptible range for all tested antimicrobials and were categorized as beta-lactam susceptible with benzylpenicillin disc screen, except, as anticipated, beta-lactamase producing isolates. Surprisingly, the same results were obtained for strains with amino acid substitutions of the group M (see **Figure 1**). Based on our crystal structure analysis, we suggest that the group I and group II substitutions Arg517His and Asn526Lys influence the binding of beta-lactam antimicrobials trough a common mechanism by disrupting the Asn526-Arg517-Glu324 interaction (see **Figure 2e**). Since these three amino acids are able connect the beta sheet and the alpha helix that harbour the KTG and STVK motifs, their interaction might be important for a proper binding site formation. Further molecular investigations including site-directed mutagenesis of these amino acids are clearly needed to test this hypothesis. Interestingly, a just recently published study on the crystal structure of *H. influenzae* comes to a similar conclusion that mutations Arg517His and Asn526Lys might lead to resistance through long-range structural rearrangements (37). Our study complements this finding and proposes a mechanistic explanation, i.e. the disruption of the Asn526-Arg517-Glu324 bridge.

Group M substitutions are distant to the binding site and shown in **Figure 4** and have probably no influence on the configuration of the binding site. We propose, that strains belonging to group M are phenotypically equal to the wild type, since an impact on the beta-lactam binding site is unlikely. This was confirmed by low MICs in antimicrobial susceptibility testing of these isolates (**Figure 1**) and categorization as beta-lactam susceptible in benzylpenicillin disc screen (**Figure S2**).

Investigation of group III alterations (even though not found in this study) based on PBP3 structure of *P. aeruginosa* was not possible due to differences in amino acid sequences between *P. aeruginosa* and *H. influenzae* in this region.

Although Ellington *et al.* concluded that prediction of antimicrobial resistance based on whole genome sequencing data is currently not feasible and requires collection of extensive data sets (38), the mechanistic understanding on how single point mutations impact binding affinities of antibiotic drugs may help to predict antimicrobial resistance in the future.

All isolates in this study showed very low MICs to piperacillin-tazobactam (range <0.016-0.25 mg/l), which was also documented in european and asian studies (14, 39). Piperacillin MICs were also very low in non-beta-lactamase-producing strains (range <0.016-0.25 mg/l), which implicates that there is no intrinsic activity of tazobactam against *Haemophilus* strains. Nevertheless, no EUCAST nor CLSI breakpoints for *H. influenzae* have been issued to date for piperacillin, and only CLSI breakpoints for piperacillin-tazobactam have been published (29).

Reliable detection of beta-lactam resistance in *H. influenzae* is essential, since beta-lactam antimicrobials represent a first line treatment for infections caused by this bacterium. Detection of *H. influenzae* strains producing beta-lactamase with nitrocefin-based tests is rapid and reliable. In 34 out of 36 isolates with PBP3 alterations, EUCAST screening for beta-lactam susceptibility with benzylpenicillin disc was able to separate between PBP3 alterations with no impact on the beta-lactam binding (group M) and the alterations leading to decreased affinity to beta-lactams (group I and II). All isolates containing amino acid substitutions of group M were categorized as beta-lactam susceptible (showing zone diameters of 15 to 19 mm), together with wild type isolates (exception: beta-lactamase producing strains) as shown in **Figure S2**. Two isolates belonging to groups I and II were falsely categorized in screening as beta-lactam susceptible, but it needs to be mentioned that they both exhibited zone diameter on the breakpoint (12 mm), and that technical factors could be responsible for this borderline zone diameter. Further studies on larger collection of clinical isolates are needed to investigate this observation. Furthermore, our study showed that some gBLNAR strains (belonging to group I and II) are categorized as susceptible for amoxicillin, ampicillin and amoxicillin-clavulanic acid when applying EUCAST interpretation criteria (**Table 4**). Although clinical impact of these in vitro findings needs some further investigations, two pragmatic approaches were recommended by Skaare et al. (35): (a) *H. influenzae* isolates with altered PBP3 positive by screening for beta-lactam resistance with benzylpenicillin 1U disc should be categorized as cefuroxime resistant and always be reported as ampicillin resistant in cases of meningitis; (b) a comment should be added recommending high-dose aminopenicillin therapy or the use of other agents in severe infections caused by screening-positive isolates categorized as susceptible to aminopenicillins by disc or gradient diffusion.

Emergence of strains with resistance to carbapenems is another worrying development. The study of Cherkaoui and colleagues provides evidence that development of imipenem heteroresistance depends on a combination of altered PBP3, slowed drug influx and its enhanced efflux due to the loss of regulation of the efflux pump (33). In the present study, we found that 15% of BLNAR strains were categorized as meropenem intermediate when interpreted with EUCAST meningitis breakpoints (**Table 4**). In contrast, meropenem resistant isolates were reported in the 7.4% of the gBLNAR isolates from a Portugese study (21). The same study showed that the MIC_50_ and MIC_90_ values of meropenem were both four times higher for gBLNAR isolates compared to gBLNAS. The same pattern was also observed for the strains of this study, but no strains with resistant phenotype were observed (**Table 4**).

Our study has the following limitations: (a) the number of isolates examined in this study is limited, but the results are in accordance with the findings of other Swiss and European studies (3, 5, 14, 17); (b) no group III or group III-like isolates were included in the analysis, since these isolates are still rarely found in Europe (3, 14, 17); (c) besides the heterogenous impact of various alterations in PBP3 with multiple changes in amino acids also other possible resistance mechanisms like efflux pumps might further act on the MICs of antimicrobials (17, 33, 40).

In conclusion, based on our molecular and phenotypic findings, we assume that strains belonging to M group are phenotypically equal to wild type, while amino acid substitutions of group I and group II affect the beta-lactam binding through a common mechanism by disrupting the Asn526-Arg517-Glu324 interaction. Not all PBP3 alterations have an influence on the resistant phenotype in *H. influenzae*. When EUCAST interpretation criteria were applied, some gBLNAR strains (all harbouring group I and II mutations) were categorized as susceptible to ampicillin, amoxicillin or amoxicillin-clavulanic acid.

## Acknowledgements

We thank Helena Seth-Smith for critically reading the manuscript.

## Compliance with Ethical Standards

### Ethical approval

This article does not contain any studies with human participants or animals performed by any of the authors.

### Informed consent

Not applicable (this article does not contain any studies with human participants by any of the authors).

## Online ressources and electronic supplementary material (ESM)

**ESM1:** Repository with raw data including all analysis scripts and figure-generating scripts in R: https://gitlab.com/csb.ethz/haemophilus-pbp3-manuscript

**ESM2:** Supplementary figures can be found at the end of this document.

**Fig. S1** Determined MIC values for all isolated strains (*bla*-positive and *bla*-negative).

**Fig. S2** Distribution of investigated *H. influenzae* strains (n=53) according to benzylpenicillin 1 U disc zone diameter (screening for beta-lactam resistance according to EUCAST).

**Fig. S3** Location of amino acid substitutions in the protein sequence of the reference strain *H. influenzae* RD KW20 (UniProt Entry P45059).

**ESM3:** Annotated sequence aligment between *H. influenzae* RD KW20 (UniProt entry P45059) and *P. aeruginosa* (UniProt entry Q51504). The file can be downloaded from the gitlab repository: https://gitlab.com/csb.ethz/haemophilus-pbp3-manuscript

## Supplementary Material

**Fig. S1.**
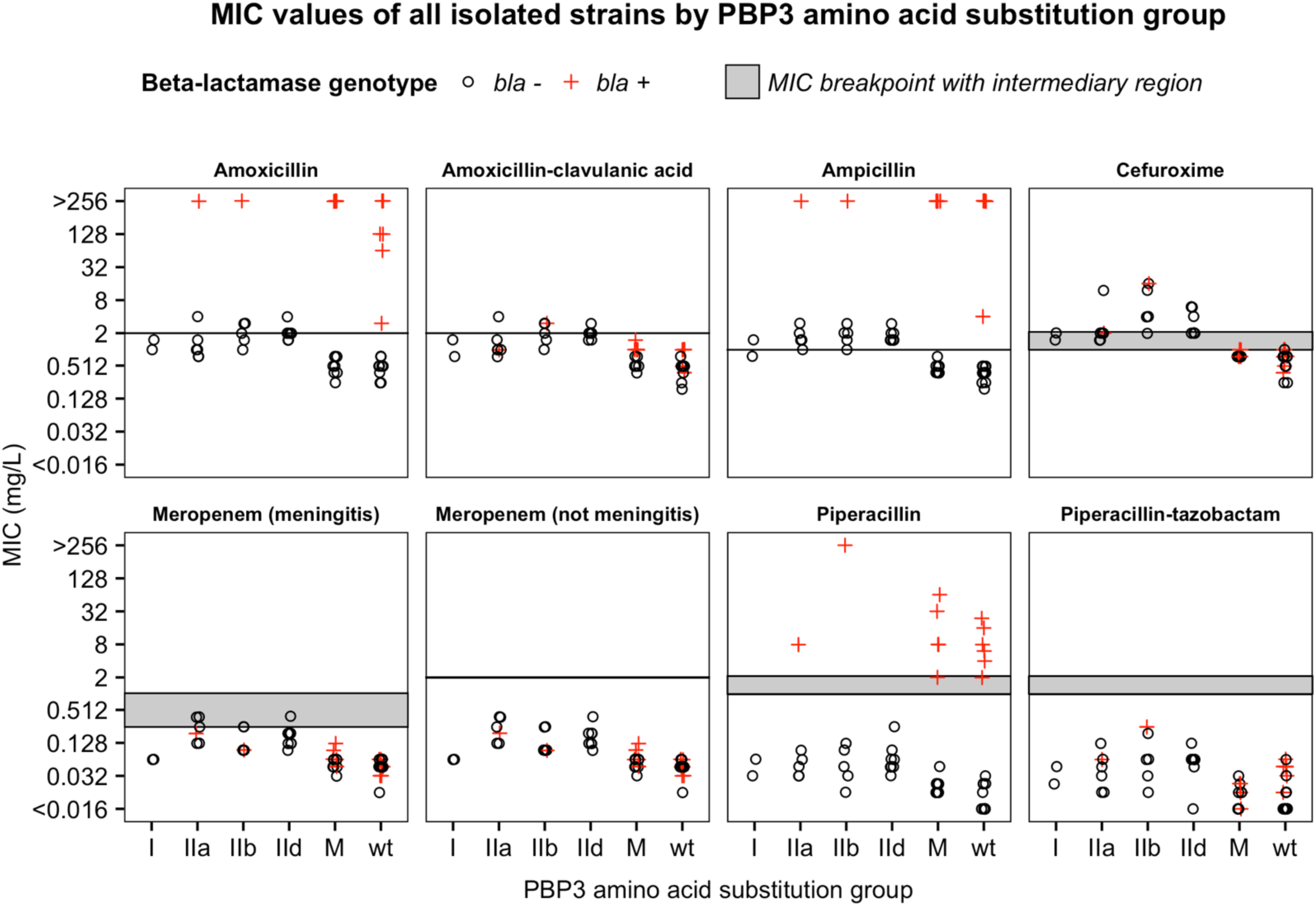
Minimal inhibitory concentrations (MIC) of all isolated strains for tested antibiotics (amoxicillin, amoxicillin-clavulanic acid, ampicillin, cefuroxime, meropenem (meningitis), meropenem (not meningitis), piperacillin, piperacillin-tazobactam). The horizontal line indicates the MIC breakpoint and, if applicable, the grey region indicates the intermediary region. All antibiotic breakpoints are according to EUCAST with the exception of piperacillin and piperacillin-tazobactam which are according to CLSI.

**Fig. S2.**
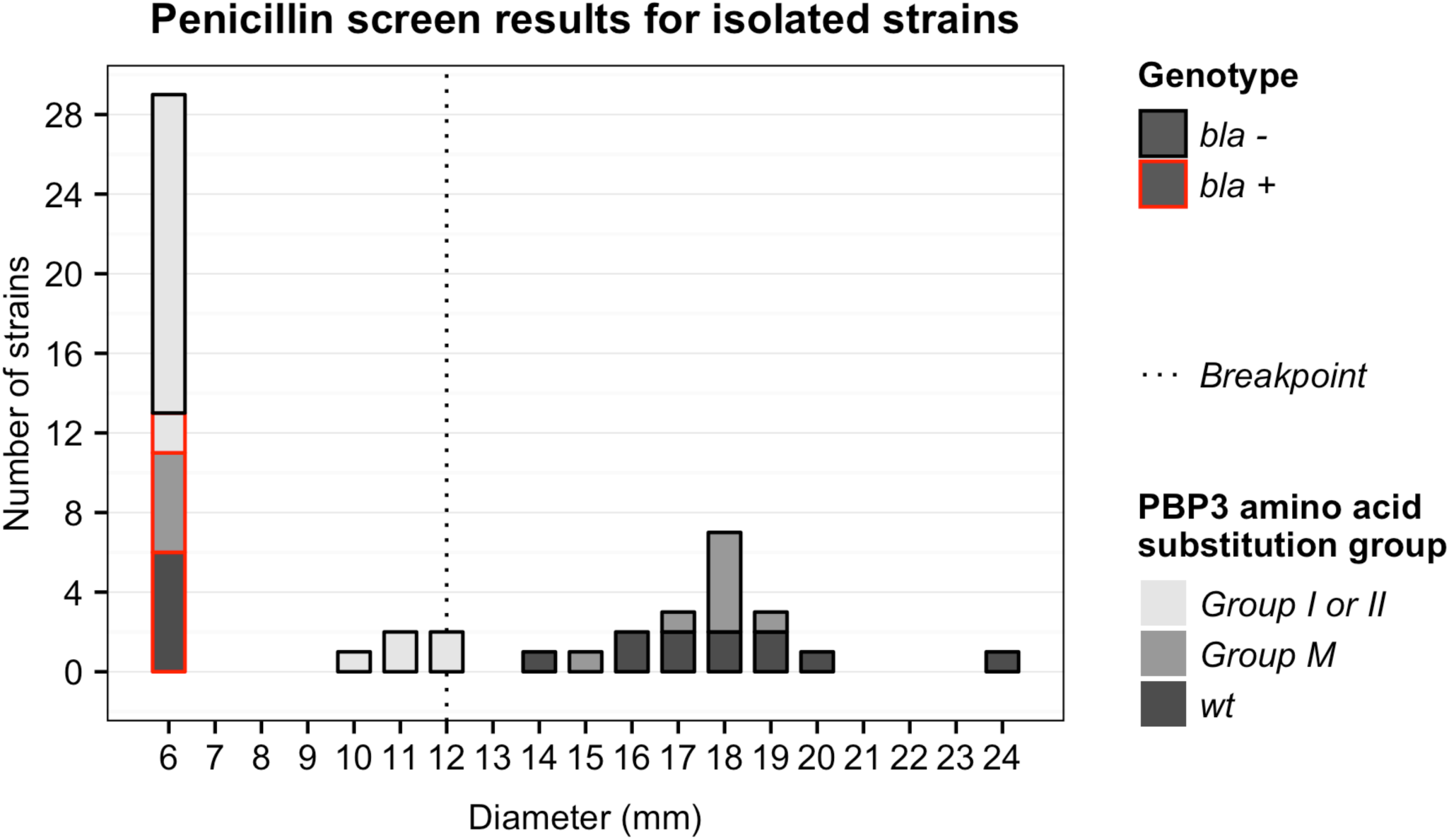
Distribution of investigated *H. influenzae* strains (n=53) according to benzylpenicillin 1 U disc zone diameter (screening for beta-lactam resistance according to EUCAST). Wildtype: n=17; group M: n=13; group I and II: n=23; beta-lactamase positive isolates (red border): n=13. The dotted line indicates the diameter breakpoint (resistant ≤11 mm, susceptible ≥12 mm).

**Fig. S3.**
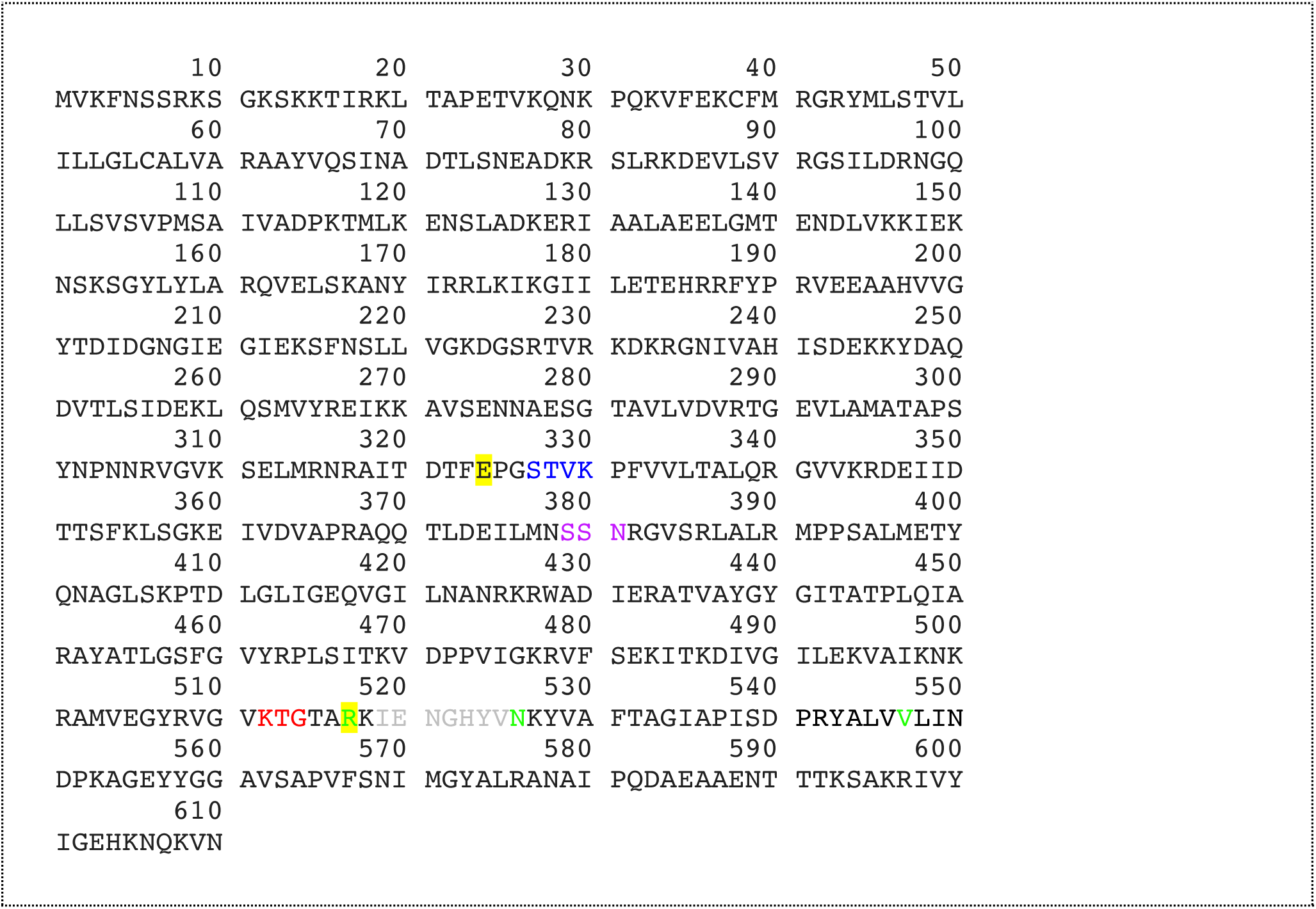
Protein sequence of the reference strain *H. influenzae* RD KW20 (UniProt Entry P45059). The group-specific amino acid substitutions Arg517His, Asn526Lys and Val547Ile are shown in green and their suggested interaction partners Arg517 and Glu324 are highlighted in yellow. The three motifs that define that active site of the serine penicillin-recognizing enzyme family are shown in blue, purple and red. The amino acid substitution Arg517His were found in all group I strains and might lead to a loss of an electrostatic interaction to Glu324. Substitution Asn526Lys were found in all group II strains and might lead to a loss of a hydrogen bond to Arg517. Substitution Val547Ile were found in all group M and in some group II strains and has most likely no impact on the protein structure. For more information on the identified mutations, see Table 1. For a representation of the mutations in the three-dimensional protein structure, see **Figures 4-6**. Amino acids in grey belong to a beta-loop that is not represented in the crystal structure that was used for structure analysis (PDB ID 3PBR).

